# Attentional Bias Modification is associated with fMRI Response towards Negative Stimuli in Residual Depression: A Randomized Controlled Trial

**DOI:** 10.1101/322842

**Authors:** E. Hilland, N.I. Landrø, C. J. Harmer, M. Browning, L. A. Maglanoc, R. Jonassen

## Abstract

**Background:** Modification of attentional biases (ABM) may lead to more adaptive emotion perception and emotion regulation. Understanding the neural basis of these effects may lead to greater precision for future treatment development. Task-related fMRI following ABM training has so far not been investigated in depression. The main aim of the RCT was to explore differences in brain activity after ABM training in response to emotional stimuli.

**Methods:** A total of 134 previously depressed individuals were randomized into 14 days of ABM- or a placebo training followed by an fMRI emotion regulation task. Depression symptoms and subjective ratings of perceived negativity during fMRI was examined between the training groups. Brain activation was explored within predefined areas (SVC) and across the whole brain. Activation in areas associated with changes in attentional biases (AB) and degree of depression was explored.

**Results:** The ABM group showed reduced activation within the amygdala and within the anterior cingulate cortex (ACC) when passively viewing negative images compared to the placebo group. No group differences were found within predefined SVC’s associated with emotion regulation strategies. Response within the temporal cortices was associated with degree of change in AB and with degree of depressive symptoms in ABM versus placebo.

**Limitations:** The findings should be replicated in other samples of depressed patients and in studies using designs that allow analyses of within-group variability from baseline to follow-up.

**Conclusions:** ABM training has an effect on brain function within circuitry associated with emotional appraisal and the generation of affective states.

*Clinicaltrials.gov identifier: NCT02931487*

## Introduction

A number of effective treatments exist for major depressive disorder. However, following successful treatment relapse is common with 50-70% of patients relapsing within 5 years.^1,2^ Residual symptoms are among the strongest predictors for relapse in recurrent depression.^3^ Cognitive theories of depression posit that biased information processing for emotional stimuli plays a key role in development and relapse in depression.^4^ Despite mixed findings, clinically depressed subjects, as well as currently euthymic previously depressed subjects, have repeatedly been reported to orient their attention toward negative faces rather than neutral or positive faces.^5–10^ Attentional biases (AB) and deficits in cognitive control may interfere with emotion regulation and mood state. Negative cognitive biases in depression are thought to be facilitated by increased influence from subcortical emotion processing regions combined with attenuated top-down cognitive control.^4,11^

Computerized ABM procedures aim to implicitly retrain biased attentional patterns.^12^ Although there is debate about the true effect size of ABM in depression^13,14^ some studies have reported reduced depressive symptoms after successful modification of AB.^15–18^ The neural basis of changes in AB, which is believed to be the mechanism of change behind symptom improvement after ABM training, has so far not been investigated. The functional neurobiology of emotion perception distinguishes between structures critical for appraisal, generation of affective states and emotion regulation. The amygdala and insular cortex are particularly important within a ventral system linked to the emotional significance of stimuli, and the production of affective states. The ventral ACC plays a main role in automatic regulation of emotional responses. A dorsal system includes the dorsal ACC and prefrontal regions and is argued to be involved in effortful regulation of affective states and subsequent behavior.^19,20^

The neural effects of a single session ABM in healthy individuals include lateral prefrontal cortex reactivity towards emotional stimuli^21^ indicating moderation of the dorsal neurocircuitry in emotion perception. One fMRI study in young women with sub-threshold depression found differences between ABM and placebo in measures of spontaneous fluctuations within the right anterior insula and right middle frontal gyrus,^22^ areas critical for emotion generation and automatic emotion regulation of emotional responses. In a study with clinically depressed participants differences in resting state connectivity between ABM and placebo was found within the middle frontal gyrus and dorsal ACC, a neural system important for cognitive control over emotions, along with changes in a network associated with sustained attention to visual information in the placebo group.^23^ Overall, these early results provide some evidence that ABM modifies function in emotional regulatory systems although the small study sample sizes and variety of approaches used may underpin the absence of consistent effects across studies.^24^

No study has investigated ABM-induced changes in emotion processing using fMRI in a large clinical sample after multiple training sessions. In this pre-registered clinical trial (NCT02931487) we used a sample of 134 participants previously treated for depression and with various degrees of residual symptoms. A main aim was to explore the neural effects of ABM within both ventral- and dorsal emotion perception circuitry. We measured BOLD response within well-established emotion regulation circuitry, based on previous studies in response to emotionally arousing stimuli when participants attempted to actively regulate their emotional response. Brain activation in response to passive viewing of negative stimuli was explored across the whole brain and within the bilateral amygdala. Furthermore, we examined how changes in AB, the mechanism by which ABM is believed to work, and changes in symptoms differed between ABM as compared to placebo.

## Methods and materials

### Participants and Screening procedures

Patients that had been treated for at least one previous episode of MDD were randomized into two treatment conditions with either a positive ABM- or a closely matched active placebo training condition. Block randomization (1:1) was performed at inclusion to ensure equal numbers of participants and similar characteristics for the two groups. Participants were invited to be part of the fMRI study immediately after training and preferably within one week after ABM training. The current clinical trial (NCT02931487) is an extension of a larger double-blinded randomized clinical trial (RCT)(NCT02658682) including 321 patients with a history of depression. A total of 136 eligible participants between 18-65 years old were enrolled for fMRI.

The main recruitment site was an outpatient clinic in the Department of Psychiatry, Diakonhjemmet Hospital in Oslo. Participants were also recruited from other clinical sites and via social media. Individuals diagnosed with current- or former neurological disorder, psychosis, bipolar spectrum disorders, substance use disorders, attention deficit disorder, or head trauma were excluded via pre-screening. Informed consent was obtained before enrolment. The procedure was approved by The Regional Ethical Committee for Medical and Health Research for Southern Norway (2014/217/REK sør-øst D).

Inclusion criteria were individuals that had experienced more than one depressive episode fulfilling the Mini International Neuropsychiatric Interview (M.I.N.I 6.0.0) A1a (depressed mood) and/or A2a (loss of interest or pleasure) criteria, more than 5 positive items on A3 and filling the A5 criterion (DSM 296.30-296.36 Recurrent/ ICD-10 F33.x). To assess both clinically- and self-rated of symptoms Beck Depression Inventory (BDI-II)^25^ and Hamilton Rating Scale for Depression (HDRS)^26^ were administered.

### Attentional bias modification procedure

The ABM task was a computerized visual dot-probe procedure developed by Browning and coworkers.^15^ A fixation cross was initially displayed followed by two images (the stimuli) presented concurrently on the top and bottom of the computer screen. Following stimulus onset, a probe (one or two dots) immediately appeared on the same location as one of the image stimuli and remained on the screen until the participant responded. The types of stimuli were pictures of emotional faces of three valences; positive (happy), neutral, or negative (angry and fearful). A single session of the task involved 96 trials with equal numbers of the three stimulus pair types. In addition, equal numbers of trials were randomly presented for 500- or 1000 ms before the probe was displayed. In each trial of the task, stimuli from two valences were displayed, in one of the following pairing types: positive-neutral, positive-negative, and negative-neutral. In the ABM condition, probes were located behind positive stimuli in 87 % of the trials (valid trials), as opposed to 13% with probes located behind the more negative stimuli (invalid trials). Consequently, participants should implicitly learn to deploy their attention toward positive stimuli, and in this way develop a more positive AB when completing the task. The neutral ABM placebo condition was otherwise identical, except the location of the probe, which was located behind the positive (valid trials) stimuli in 50% of the trials. Participants completed two sessions (96 trials) of ABM daily during the course of fourteen days (28 sessions in total) on identical notebook computers (14″ HP EliteBook 840, 1600×900, 8GB, Intel Core i5-4310U), which were set up and used exclusively for ABM-training.

### MRI Scan acquisition

Scanning was conducted on a 3T Philips Ingenia whole-body scanner, with a 32 channel Philips SENSE head coil (Philips Medical Systems). Functional images were obtained with a single-shot T2* weighted echo planar imaging sequence (repetition time (TR): 2000 ms; slice echo time (TE): 30 ms; field of view (FOV): 240×240×117; imaging matrix: 80×80; flip angle 90°, 39 axial slices, interleaved at 3 mm thickness, no gap, voxel size 3×3×3 mm). The scanning session consisted of 340 volumes, synchronized to the onset of the experiment. Slice orientation was adjusted to the line running from the anterior to posterior commissure. A T1-weighted anatomical image with a voxel size of 1×1×1 mm was recorded for registration of the functional images (TR: 8.5 ms; TE: 2.3 ms; FOV: 256×256×184; flip angle: 7°;184 sagittal slices).

### fMRI Experimental procedure

The study used a modified emotion regulation experiment. Participants were scanned as they were viewing sequences of negative and neutral images while carrying out instructions either to down-regulate their emotional responses using a reappraisal strategy, or to simply allow themselves to attend to the pictures without trying to influence their emotional reactions. After each image the participants provided a rating of the intensity of their emotional state using a visual analogue scale (VAS) ranging from neutral to negative. Stimuli were selected from the International Affective Picture System ^27^ and the Emotional Picture Set.^28^ Negative and positive pictures were counterbalanced concerning their normative valence and arousal ratings (see Supplemental information for more detail). Each trial started with a fixation cross followed by a written instruction, (“Attend” or “Regulate”). The instruction was presented for 2000 ms. A negative or neutral image was presented for 6000 ms, followed by a rating screen time-locked to 6000 ms. Between stimuli there was a temporal jitter randomized from 2000-8000 ms (mean ISI; 3,700 ms) to optimize statistical efficiency in the event related design.^29^ The task consisted of blocks of 18 trials with a 20 second null-trial between the two blocks. The procedure was completed in two independent runs during the scanning session, 72 trials in total. In each block 12 items were neutral and 24 items were negative, giving three counterbalanced experimental conditions; AttendNeutral, AttendNegative, RegulateNegative. The stimulus-order in each block was interspersed pseudo-randomly from 12 unique lists. The total duration of one single functional scanning run was ~11 minutes, and total scan time~22 minutes. Stimuli were presented using E-Prime 2.0 software (Psychology Software Tools). An MRI compatible monitor for fMRI was placed at the end of the scanner behind the participants’ head. Participants watched the monitor using a mirror placed at the head coil. Responses were collected with a response grip with two response buttons. Physiological data (heart and respiration curves) were recorded at 1000 Hz using a clinical monitoring unit digitized together with scanner pulses.

### Training and instruction procedures

A written protocol with detailed instructions was used to introduce the emotion regulation experiment. The protocol was dictated for each participant by the researcher outside the MRI-scanner in order to standardize the verbal instructions. The fMRI experiment had three in-scanner exercise trials before scan start in order to make participants familiar with the instructions, timing, response buttons and VAS scale. The training procedure was repeated before the second run of the experiment.

### Symptom change and subjective ratings of negativity

Changes in self-rated and clinician rated symptoms were analyzed in PASW 25.0 (IBM) using a repeated measures ANOVA with intervention (ABM versus placebo training) as a fixed factor. Symptoms at baseline and at two weeks follow-up (time) were the dependent variable. To investigate self-reported emotional reactivity (VAS scores) during fMRI a factor based on the three experimental conditions AttendNeutral, AttendNegative, RegulateNegative was added as factor and analyzed in a repeated measures ANOVA.

### fMRI analyses

Whole brain analysis used the AttendNegative > AttendNeutral contrast to tests whether ABM influences overall brain activity in response passive viewing of emotional stimuli. Clinician-rated (HRSD) symptoms at baseline was demeaned and used as covariate. Spatial smoothing FWHM was set to 5 mm. Featquery was used for FEAT result interrogation. Mean local percent signal change was extracted to explore individual distribution within significant clusters from FEAT. Interaction analysis was performed in order to test whether areas within the brain respond differently in ABM versus placebo in relation to AB and symptom change.

Small volume correction (SVC’s) used regions from a recent meta-analysis on neuroimaging and emotion regulation.^30^ This meta-analysis is comprised of 48 neuroimaging studies of reappraisal where the majority of studies involved downregulation of negative affect. Buhle et al^30^ reported seven clusters related to emotion regulation consistently found within prefrontal cognitive control areas when contrasted to passive viewing of negative images. The clusters were situated in left and right middle frontal gyrus, right inferior frontal gyrus, right medial frontal gyrus, left and right superior temporal lobe, and left middle temporal gyrus (See Supplemental). The bilateral amygdala, but no other brain regions was reported for the opposite contrast comparing negative viewing to emotion regulation. Brain activation derived from passive viewing of negative images as compared to passive viewing of neutral images was not included in the results from the meta-analysis.^30^

Binary spheres with a 5 mm radius based on MNI coordinates of peak voxels were used for the predefined regions. Two single masks were created for the emotion regulation contrast (RegulateNegative > AttendNegative, AttendNegative > RegulateNegative). The seven cortical spheres and the two subcortical spheres respectively were combined into two single binary SVC’s. Z-threshold was set to 2.3 and cluster p-threshold was .05. Mean local percent signal change was extracted from the two SVC’s to explore individual distribution within significant clusters from FEAT. Again, clinician-rated (HRSD) symptoms at baseline was demeaned and used as covariate also in the SVC analysis.

### fMRI data preprocessing and noise reduction

The FMRIB Software Library version (FSL version 6.00) (www.fmrib.ox.ac.uk/fsl)^31,32^ was used to pre-process and analyze fMRI data. FMRI data processing was carried out using FEAT (FMRI Expert Analysis Tool) Version 6.00, part of FSL (FMRIB’s Software Library, www.fmrib.ox.ac.uk/fsl). In conjunction with FEAT FSL-PNM, 34 EVs were applied to regress out physiological noise from pulse and respiration.^33^ Registration to high resolution structural and/or standard space images was carried out using FLIRT.^34,35^ Registration from high resolution structural to standard space was then further refined using FNIRT nonlinear registration.^36^ All registrations were manually inspected to ensure proper alignment. Time-series statistical analysis was carried out using FILM with local autocorrelation correction.^37^ Linear registration was conducted with 12 DOF. Z (Gaussianised T/F) statistic images were thresholded using clusters determined by Z>2.3 and a (corrected) cluster significance threshold of P=0.05.^38^ Two participants were excluded from the analyses due to signal loss caused by a technical problem with the head coil. Time series from each subjects’ two first level runs were combined using an intermediate fixed effect model in FEAT before submission to second level analysis. A total of 134 participants, 64 from the ABM group and 70 from the placebo group were included in the intermediate - and the higher level FEAT analysis at group level.

## Results

### Sample characteristics

**Table 1.** shows means, standard deviations (SD) or numbers (%).ISCED= International Standard Classification of Education. SSRI= any current usage of an antidepressant belonging to the Selective Serotonin Reuptake Inhibitors MDE=Major Depressive Episodes according to M.I.N.I.

|  | Placebo (n=70) | ABM (n=64) |
| --- | --- | --- |
| **Age** | 39,2 (13.4) | 39,0 (12.8) |
| **Gender (females)** | 44 (62) | 47 (73) |
| **Education Level (ISCED)** | 5,9 (1.2) | 5,8 (1.2) |
| **Medication (SSRI)** | 23 (33) | 22 (34) |
| **Number of previous MDE** | 4,4 (5.3) | 4,4 (7.6) |
| **Days between ABM and fMRI** | 6.9 (8.7) | 6.6 (7.2) |
| ***Baseline symptoms:*** |  |  |
| **HRSD** | 7,6 (4.7) | 9,3 (6.1) |
| **BDI-II** | 12,1 (8.7) | 17,12 (11.6) |

### Symptom change after ABM

There was a statistically significant effect of the intervention (time) for rater-evaluated depression as measured by the change in HRSD, with lower symptoms of depression in the ABM group [F (1,132) = 4.277, η ^2^= .03, p = .041]. The means and standard deviations at baseline in ABM was (9.56 (6.38)) and placebo (7.53 (4.69)) and changed to (7.93 (5.90)) and (7.77 (5.76)) at two weeks follow-up. No statistically significant effect was found for self-reported symptoms as measured by the BDI-II [F (1,132) = 2.048, p = .155]. The means and standard deviations at baseline in ABM was (17.12 (11.62)) and placebo (12.09 (8.66)) and changed to (13.25 (12.04)) and (9.82 (8.72)) at two weeks follow-up. There was a general symptom improvement in both the ABM and placebo group as measured by the BDI-II from baseline to post training [F (1,132) = 29.775, η^2^= .18, p < .001]. This is in accordance with the results from the sample in which this smaller cohort of subjects is drawn (41).

### Subjective ratings of perceived negativity

We found a statistically significant difference between self-reported emotional reactivity measured by VAS scores during the fMRI experiment between task conditions. The repeated measures ANOVA showed that mean VAS scores were lowest when viewing neutral images (M=8.2(7.8)) followed by when patients were encouraged to regulate negative experience towards negative images (M=40.8 (16.9)), and highest for the passive viewing of negative images (62.0(15.3)) [F= (1,133) = .074, η^2^= .93, p < .001]. A post hoc test showed that the differences between the passive and regulate viewing conditions for negative stimuli was large and statistically significant F (1,133) = 202.81, η^2^= .60, p < .001]. VAS ratings did not differ between ABM and placebo [F= (1,133) = .993, p < .646].

### Effects of ABM from whole brain analyses

Passive viewing of negative images revealed greater placebo activation in a cluster within the pregenual ACC, the paracingulate – and the medial cortex bilaterally, extending to the right frontal orbital cortex and the frontal pole compared to ABM. The peak activation for the cluster was found in the left frontal medial cortex (MNI coordinates x y z =−16 36 −10, Z=3.86, p=.001) (Figure 1.)

**Figure 1.**
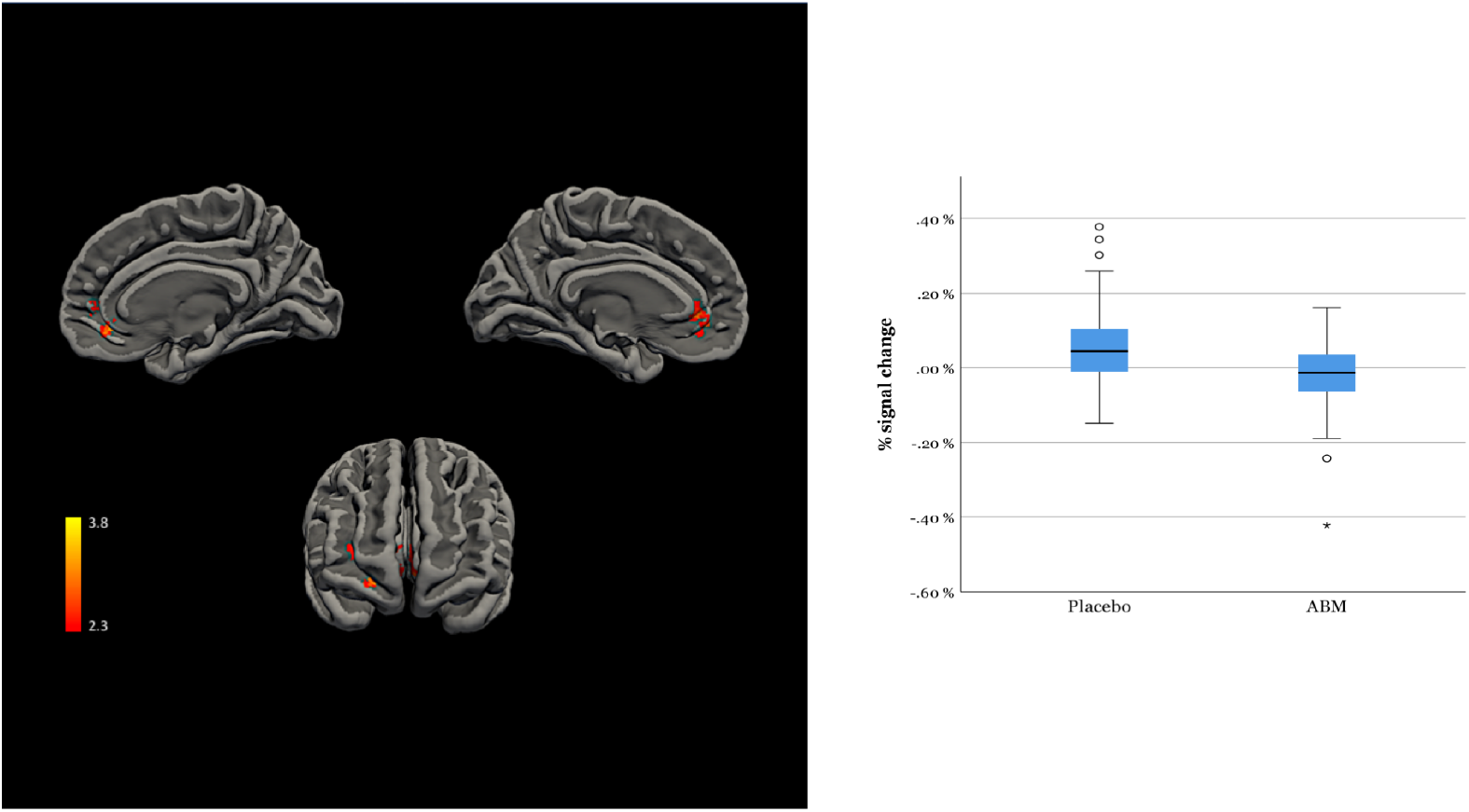
shows cluster activation (Z>2.3) for Placebo over ABM for negative images (left), together with distribution of individual percentage signal changes over significant clusters (right).

### Effects of ABM within predefined SVCs

Analyses masked across predefined emotion regulation circuitry revealed more activation placebo as compared to ABM within the right- (MNI x y z = −18 −6 −20, size=8, Z=2.89, p=.032) and left amygdala (MNI x y z = 28 0 −16, size= 3, Z = 2.55, p= .040) for the passive viewing contrast (AttendNegative > AttendNeutral) (Figure 2.).

**Figure 2.**
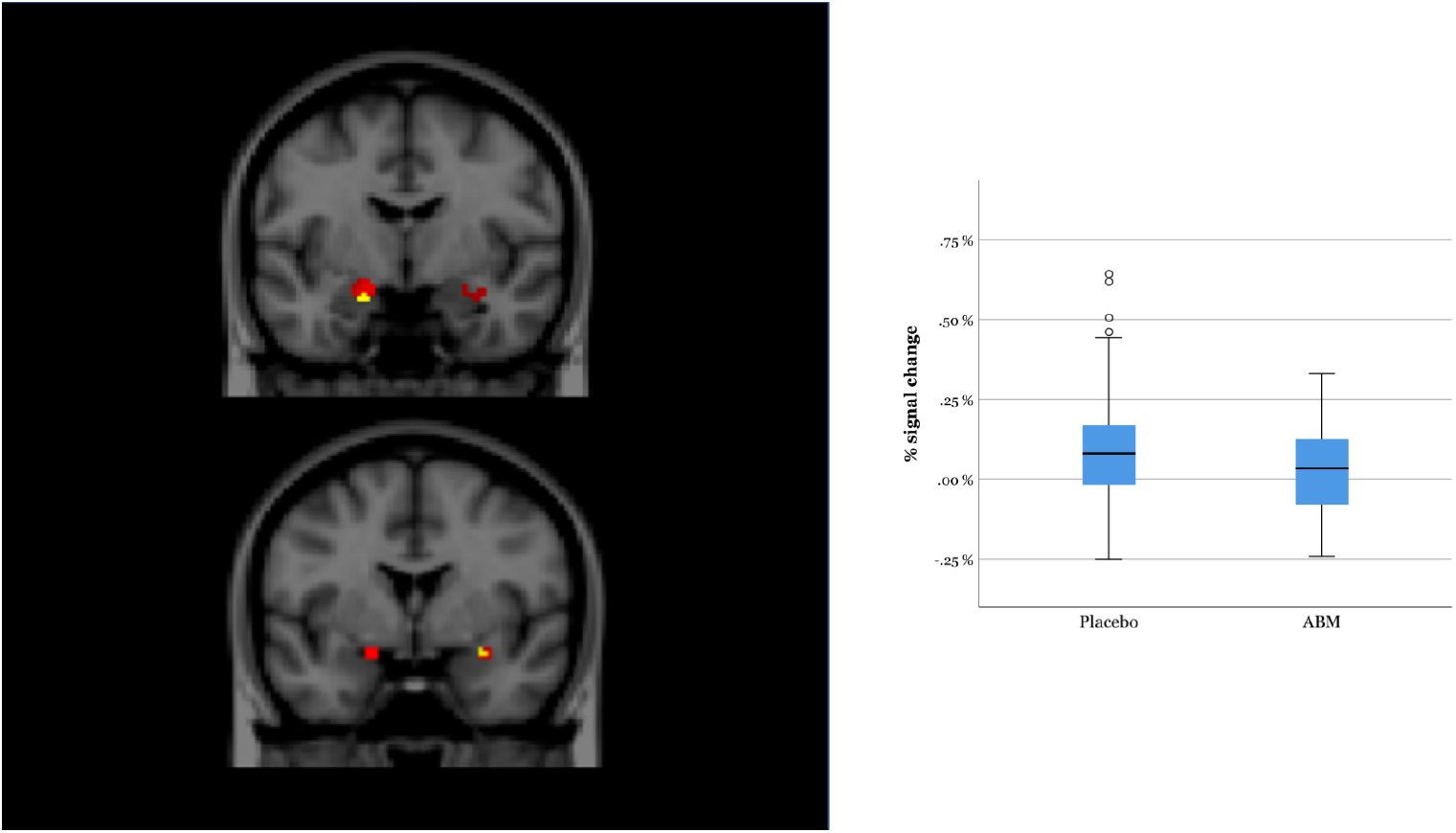
shows amygdala mean activation (left) for placebo and ABM (red) and peak voxels where placebo had more activation compared to ABM for negative images (yellow) and distribution of individual percentage signal changes over significant clusters (right).

No differences between ABM and placebo was found for the emotion regulation contrasts. Across both groups the regulate contrast (RegulateNegative > AttendNegative) revealed activation within two SVC’s in the left inferior frontal gyrus (MNI x y z = −30 −2 −54, size=75, Z=10.5, p=.017) and right middle frontal gyrus (MNI x y z = 60 26 6, size=29, Z=5.48, p=.048). The opposite contrast (AttendNegative > RegulateNegative) revealed increased bilateral amygdala activation in both the ABM- and placebo group. The largest cluster was found within the left amygdala (MNI x y z = −18 0 −14, size=75, Z=7.93; p=.006) and a smaller cluster was found within the right amygdala (MNI x y z = 26, −2, −16, size=13, Z=4.05, p=.026) (supplemental Figure1).

### Interaction with degree of attentional biases and symptom change

Two distinct clusters were associated with the interaction between the passive viewing of negative images (AttendNegative > AttendNeutral), the intervention and degree of AB change (MNI x y z = 54 −24 8, size=1061; Z=4.05, p<.001) and (MNI x y z =-50 0 10, size=547, Z=3.44; p<.020).

**Figure 3.**
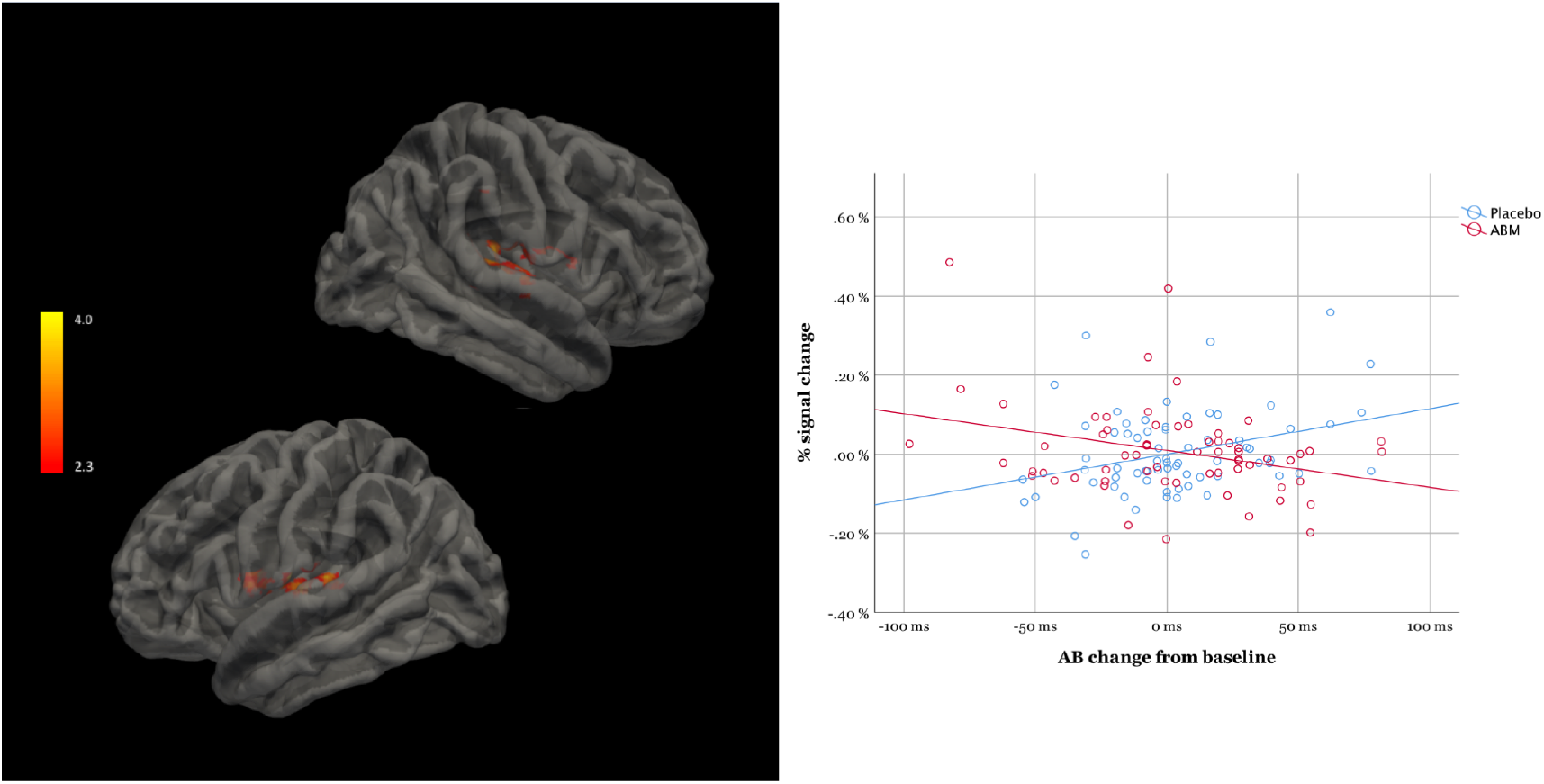
shows areas activated in association to the interaction between AB and the intervention (left). The scatter plot (right) shows the regression lines and individual distribution in the ABM and the placebo condition.

An interaction between passive viewing (AttendNegative > AttendNeutral), the intervention, and degree of symptom change (HRSD) were found within the right planum temporale and insular cortex (MNI x y z = 50 −10 18, size=872; Z=5.28, p<.001) (Figure 4.).

**Figure 4.**
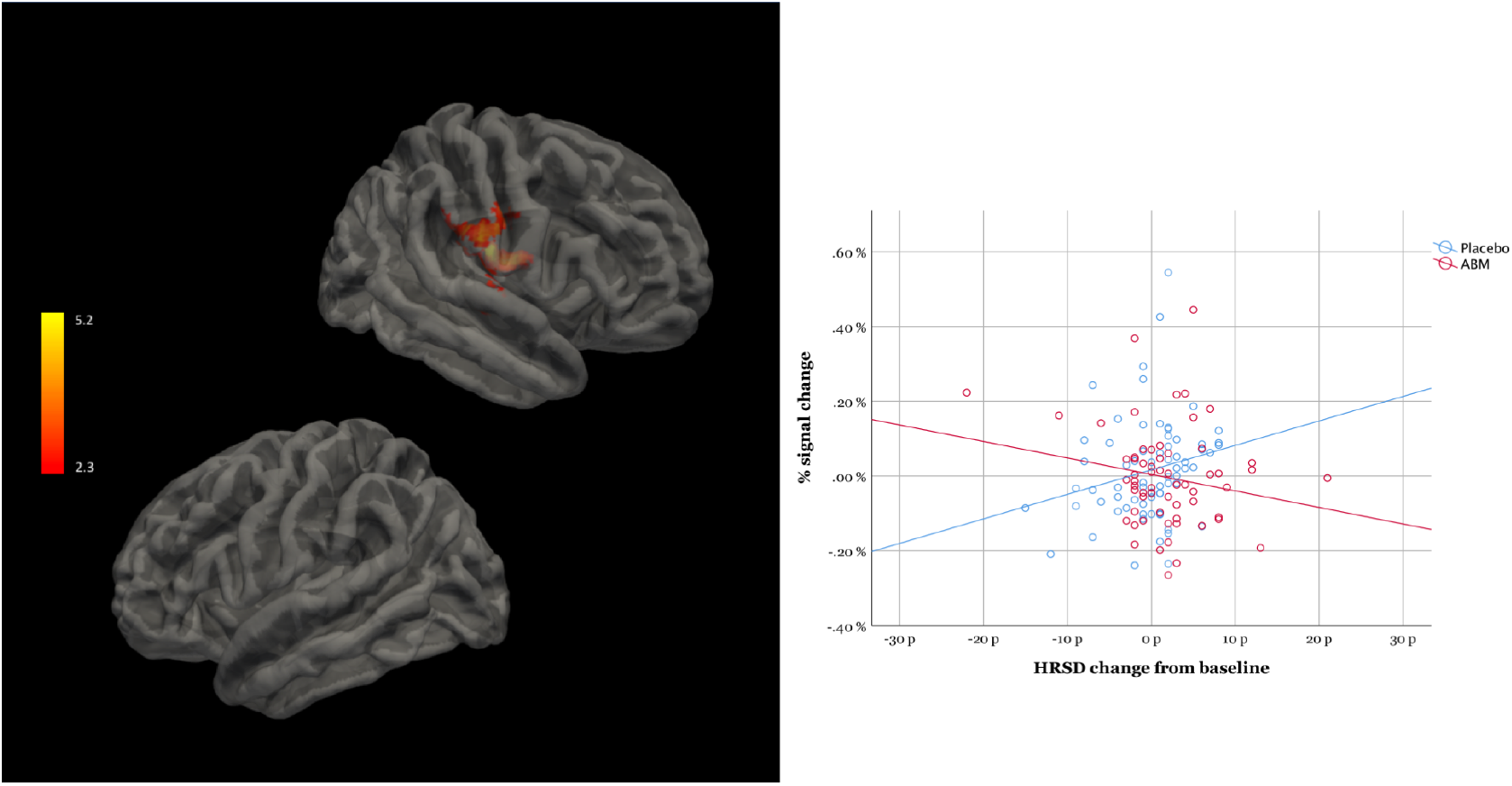
shows areas activated in association to the interaction between HRSD and the intervention (left). The scatter plot (right) shows the regression lines and individual distribution in the ABM and the placebo condition.

## Discussion

Our results revealed intervention dependent fMRI changes after two weeks of ABM within areas consistently associated with emotional appraisal and the generation of affective states, areas within a circuitry known to be altered in depression.^4,19,20^ The placebo group showed more pronounced activation for negative images within the amygdala, in midline structures, and in the pregenual ACC. Analysis of the mechanism of change showed that degree of changes in AB was linearly linked to activity in the insular cortex bilaterally. Symptom improvement after ABM was linearly associated with activation in the right insular cortex, areas involved in the generation of affective states.^19^

Analyses within predefined areas associated with effortful emotion regulation revealed activation within the left inferior frontal gyrus and right middle frontal gyrus across groups.^30^ The amygdala was also more activated during passive viewing versus active regulation of negative stimuli, but did not differ between the ABM and placebo group. In line with these results, we found no differences between ABM and placebo training as measured by subjective ratings (VAS) of perceived negativity. This is consistent with the primary outcomes from the clinical trial that found an ABM effect restricted to blinded clinician-rated, but not self-rated symptoms.^39^ Together, the results may imply that the early effects of ABM are restricted to changes in emotion generation and appraisal, as opposed to more conscious forms of emotion regulation linked to the dorsal system.

A considerable number of meta-analyses using functional connectivity in depression have shown altered activity in areas that distinguished ABM and placebo in the current study including the insula and ACC.^40–42^ In a study by Horn et al^43^ increased connectivity between pregenual ACC and insula was found in severely depressed patients compared to mildly depressed patients and healthy controls. Functional connectivity in the insula has been associated with abnormal interoceptive activity^44^ and fronto-insular connectivity has been linked to maladaptive rumination^45^ in depression. Midline brain structures including the pregenual ACC has been linked to self-referential processing,^46^ hopelessness,^47^ anhedonia^48^ and impaired emotion processing^49^ as well as with in studies of functional connectivity and depression. Notably, the ACC and insula are together with the amygdala core areas of the salience network, which determines the significance of external stimuli. The salience network has been hypothesized to play a role in switching between task positive- and negative networks^50,51^ and may well play a role in symptom improvement after ABM as found in this study.

The insula and the amygdala are among core brain areas that respond preferentially to negative stimuli in healthy individuals, and activation in the insula and ACC has repeatedly been reported across a range of experiments using emotional tasks with cognitive demand and mental imagery.^52,53^ Neural responses to negative stimuli within the amygdala, insula and ACC are found to be more pronounced in depressed patients versus healthy controls.^54^ Ma^55^ describes an emotional circuit including the insula, bilateral amygdala and ACC affected by antidepressant medication by decreasing activity towards negative- and increasing activity towards positive stimuli. Antidepressants have been hypothesized to work by remediating negative affective biases, i.e. targeting the same mechanism as when applying an ABM procedure.^56–58^ Similarly, the moderation of awareness towards negative stimuli via ABM (the mechanism of change) may alter automatic emotional vigilance and arousal towards negative stimuli. These moderations may lead to altered parasympathetic responses via circuitry involving the amygdala and ACC. The translation of these changes into improved subjective mood may take some time as the individual learns to respond to this new and more positive social and emotional perspective of the world. However, neural correlates of early changes in the processing of emotional stimuli might be a marker of a process leading to symptom improvement. This model is consistent with cognitive theories of depression^4,59^ which the ABM procedure builds on. Accordingly, studies on cognitive behavioral therapy (CBT) shows that pregenual ACC is positively correlated with the degree of symptom improvement.^60–65^ Moreover, given that that the pregenual ACC is believed to play an important role in downregulation of limbic hyperreactivity^20,66,67^ the group difference found in this study may reflect more adaptive emotion processing after ABM.

Worldwide, there is a pressing demand for evidence-based treatments in mental health. It has been argued that psychotherapy research does not provide explanations for how or why even the most commonly used interventions produce change.^68^ In a recent statement from the *Lancet Psychiatry’s Commission on treatments research in tomorrow’s science* the authors argue that there is an acute need to improve treatment and thus clinical trials should focus not only on efficacy, but also on identification of the underlying mechanisms through which treatments operate.^69^ The current study is addressing such mechanisms by targeting changes in AB, which is believed to be the mechanism, that translate into symptom improvement after ABM.

The current study is based on an RCT with a larger sample of patients that found an ABM effect on clinician-rated symptoms. It uses a well validated emotion perception task and follows a stringent pre-registered research protocol which represents a strength. This study exploits the link between a psychological mechanism, clinical measures and underlying brain function measured by fMRI, thus the results should have translational potential. The current trial is the largest study that has investigated changes in emotion processing using fMRI after ABM training.

## Limitations

A key limitation related to the research design is that fMRI assessment after ABM does not allow statistical modeling of within-individual variance from baseline to follow-up. There is an unexpected difference in symptom degrees at baseline that could be associated with group differences in brain activation. Adding symptom degree as a covariate in the fMRI analysis will regress out variance related to this particular variable. The sample consist of patients with previous depression and various degrees of residual symptoms, and needs to be replicated in studies with other patient groups. Brain activation related to ABM may also be conditionally mediated by multiple biological- and environmental factors outside the scope of this study.

## Conclusion

This study demonstrates alterations in brain circuitry linked to passive viewing, but not conscious regulation of emotional stimuli and represent the first experimental evidence of an ABM effect using task fMRI.

## Acknowledgements

We want to thank our fMRI research assistant Dani Beck. Further, we thank Diakonhjemmet Hospital, Division of Psychiatry, for help and support during the recruiting period and the Intervention Center, OUS for radiological assistance in MRI protocols, data acquisitions and screening for unexpected neuropathological findings. We thank Tor Endestad for establishing invaluable infrastructure for MRI research in our department.

We also thank our external recruitment sites; Unicare, Coperiosenteret AS, Torgny Syrstad, MD, Synergi Helse AS and Lovisenberg Hospital. The project is supported by the South-Eastern Norway Regional Health Authority, grant number: 2015052 (to NIL), Research Council Norway, grant number: 229135 (to NIL) and Department of Psychology, University of Oslo.

